# BactoMate: an integrated platform for reproducible bacterial microscopy analysis

**DOI:** 10.64898/2026.08.06.743177

**Authors:** Lasse Hallenga, Sarah Fornoff, Matthias Pesch, Dietrich Kohlheyer, Shazeb Ahmad, Jens Hör, Marc Erhardt, Philipp F. Popp

## Abstract

Quantitative microscopy of microorganisms increasingly produces large, multidimensional datasets, yet their analysis often depends on fragmented workflows spanning file conversion, segmentation, quality control, fluorescence quantification, tracking, and visualization. Here, we present BactoMate, an open-source, cross-platform graphical user interface that integrates these steps into a unified workflow for microbial image analysis. BactoMate incorporates established segmentation methods and supports both single-file and batch processing. Its modules enable image preprocessing, cell segmentation, morphology-based quality control, fluorescence and foci quantification, single-cell tracking, lineage reconstruction, structured data export, and generation of quality-control and visualization outputs. We demonstrate the applicability of BactoMate across multichannel fluorescence imaging, bacterial swimming assays, microcolony lineage analysis, phage infection assay and a microfluidic time series. All user-configurable parameters are exposed through the interface, are recorded alongside structured outputs and can be loaded for reproducible image analyses across experiments to reduce introduction of bias. By reducing workflow handoffs while preserving parameter control and exportable results, BactoMate enables accessible, reproducible, and scalable quantitative analysis of microbial microscopy data.

## Introduction

Quantitative microscopy provides a direct link between genotype, environmental conditions, and phenotypes that are difficult to resolve using population-level assays [1– 3]. At single-cell resolution, microscopy can reveal heterogeneity in cell morphology, growth and division dynamics, protein localization, intracellular complex formation, and motility. Advances in automated microscopy, microfluidics, fluorescent reporters, and camera technology now enable laboratories to acquire large, multidimensional datasets across numerous strains, conditions, fields of view, and time points [4–9]. Translating these datasets into biological insight, however, requires multiple computational steps that must be applied consistently and remain traceable throughout the analysis [10]. Image segmentation is a particularly important bottleneck. Classical approaches based on thresholding, edge detection, or watershed algorithms can perform well under controlled conditions but often require extensive tuning and manual correction when illumination, contrast, magnification, cell density, or morphology varies [11–14]. Machine-learning methods such as Omnipose [15], Cellpose [16,17] and U-NET [18] have substantially expanded the range of microbial morphologies and imaging conditions that can be analyzed, while specialized tools such as Spotiflow [19] and Trackastra [20] provide solutions for foci detection and object tracking, respectively [21]. Nevertheless, these methods are frequently operated as separate software packages or command-line tools. Connecting them to file conversion, preprocessing, morphology-based quality control, fluorescence quantification, visualization, and batch processing can therefore require custom scripts and repeated transfer of data between scripts. Existing platforms such as MicrobeJ [22] address important components of microbial image analysis but incorporating recent segmentation and tracking methods into a transparent end-to-end workflow remains challenging. Such workflow fragmentation can result in loss of metadata during analysis, undocumented parameter choices, inconsistent processing of biological replicates, and reduced reproducibility.

To address these challenges, we developed BactoMate, an open-source platform with graphical user interface that integrates microscopy file conversion, preprocessing, segmentation, morphology-based filtering, feature extraction, fluorescence analysis, foci detection, tracking, model training, data export, and visualization. Rather than replacing existing image-analysis algorithms, BactoMate connects established methods through a common interface and standardized output structure (Fig. 1). Analyses are organized within project folders in which existing image stacks, segmentation masks, parameter records, and quantitative outputs can be identified and reused. User-configurable settings are exposed in the graphical user interface and recorded in structured session logs, while persistent cell identifiers connect segmentation results to downstream measurements and trajectories. Individual modules can be used independently or combined into complete workflows for single-file and batch analysis.

**Fig. 1.**
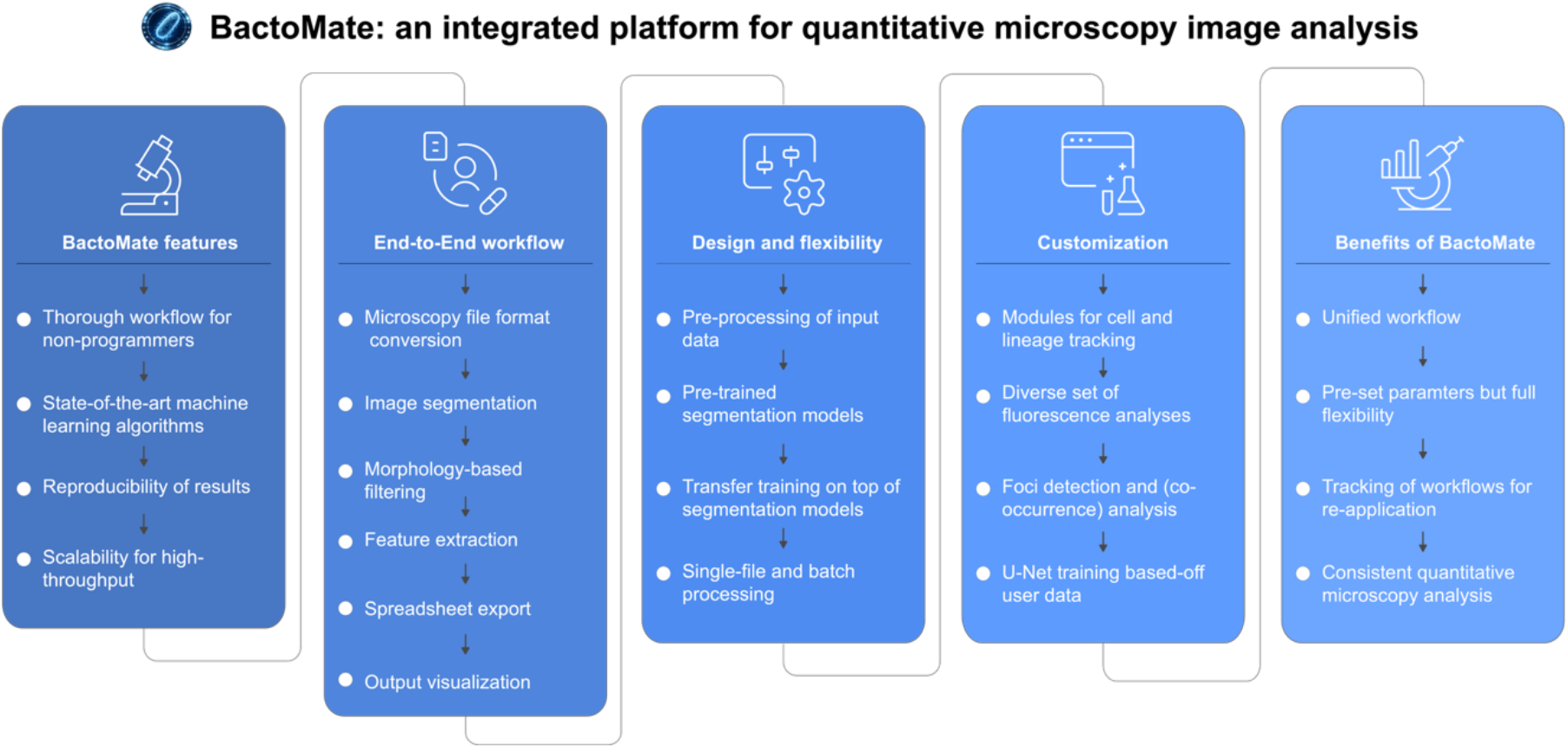
BactoMate features and workflow. (A) Schematic overview of the BactoMate workflow and its main capabilities. The software guides users from microscopy file-format conversion through image segmentation, morphology-based filtering, feature extraction, spreadsheet export, and output visualization. Additional modules support preprocessing of input data, cell and lineage tracking, fluorescence analysis, foci detection, co-occurrence analysis, model training on user-annotated data and batch processing.

Here, we describe the design and principal modules of BactoMate and demonstrate its application to diverse bacterial microscopy experiments. These include multichannel fluorescence and foci quantification, tracking of free-swimming bacteria, reconstruction of microcolony lineages, and large-scale analysis of growth under phage infection and morphology in microfluidic chambers. We distribute BactoMate through a public available GitHub repository and provide an accompanying series of tutorial videos on YouTube. Together, these examples illustrate BactoMate can support distinct biological questions while maintaining visual quality control, structured data export, and transparent parameter provenance.

## Results

### BactoMate integrates bacterial microscopy analysis into linked modules

BactoMate was developed to reduce the handoffs between separate tools used for quantitative microscopy analysis of microorganisms. The platform organizes file conversion, preprocessing, segmentation, quality control, feature extraction, tracking, visualization, and model training into linked modules within a single graphical user interface (Fig. 2). Standardized image files, segmentation masks, metadata, cell identifiers, parameter records, and quantitative outputs can be passed between modules, allowing users to follow an experiment from raw microscopy data to downstream measurements without leaving the graphical environment. Individual modules can also be used independently when only a specific analysis step is required. BactoMate analyses are organized within project folders. When a project is opened, the software identifies compatible image stacks, masks, and existing analysis outputs and suggests them as inputs for subsequent processing steps. User-configurable parameters are displayed in the interface and stored in session-specific JSON records. This project-based structure facilitates the consistent application of analysis settings across related datasets and preserves the relationship between input images, processing parameters, and exported results.

**Fig. 2.**
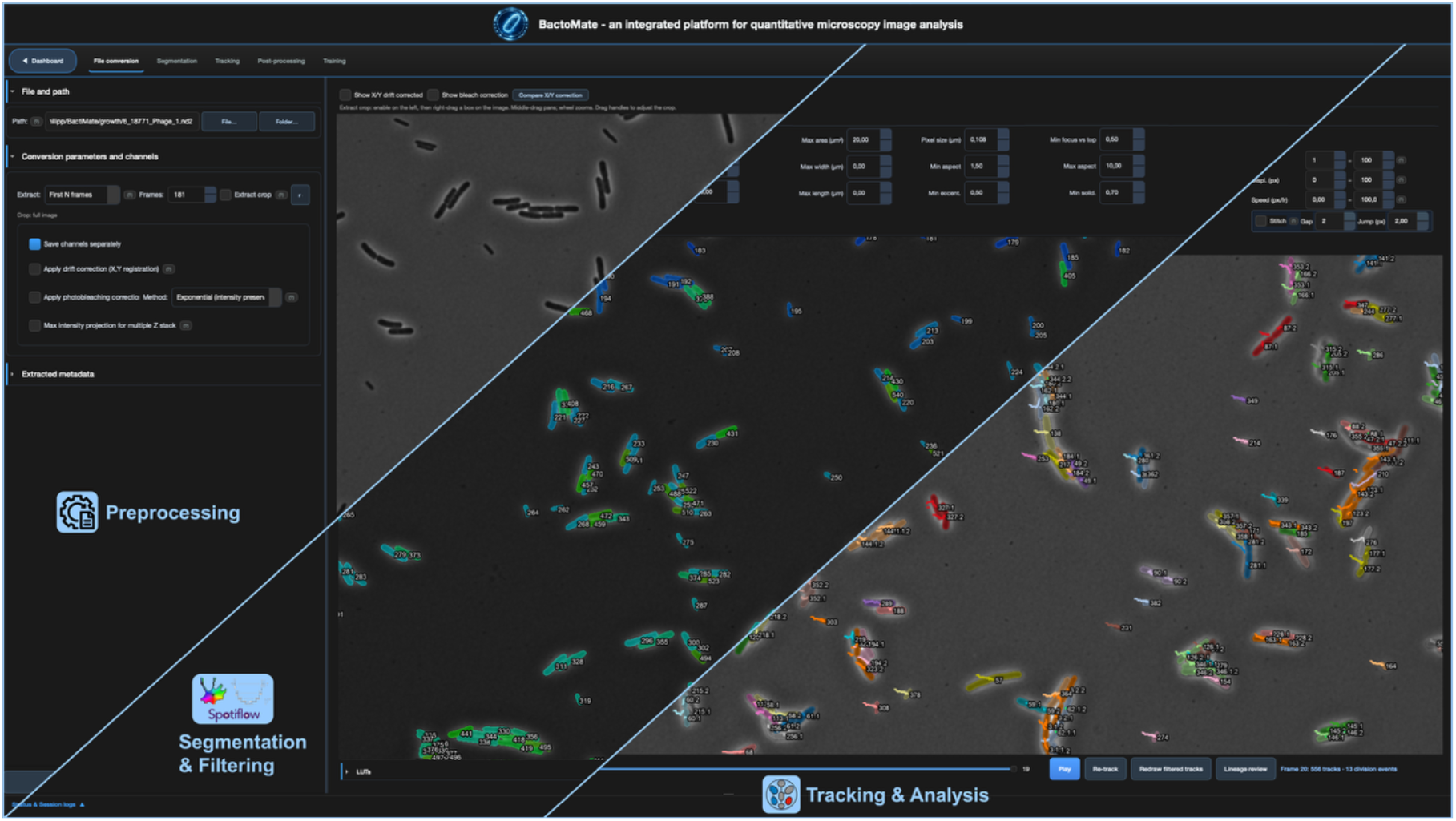
Graphical user interface of BactoMate. Left: file conversion and preprocessing controls, including image selection, cropping, drift correction, photobleaching correction, and maximum-intensity z-projection. Middle: cell segmentation and foci detection with morphology- and intensity-based filtering, distinguishing retained and excluded cells. Right: cell tracking, lineage reconstruction, and time-lapse analysis, showing color-coded trajectories, track identifiers, linking parameters, playback, manual correction, and lineage-review controls.

### File conversion, preprocessing, and image preview

BactoMate reads common proprietary and open microscopy file formats through BioIO [23] and format-specific Python libraries. Multichannel and time-resolved datasets can be inspected in an interactive preview that supports channel role assignment, selection and adjustment of lookup-table display ranges. Optional preprocessing functions include drift correction and photobleaching correction. Microscopy acquisitions are converted to multipage TIFF files for downstream analysis. Where available, pixel calibration and spatial metadata are transferred to the TIFF metadata and an accompanying JSON file. This allows measurements such as cell length and area to be reported in physical units rather than pixels and preserves essential acquisition information independently of the original proprietary file format.

### Cell segmentation, quality control, and quantitative analysis

The segmentation and analysis module connects image segmentation directly to morphology-based quality control and single-cell quantification. Users can select among integrated segmentation backends, including pretrained Omnipose models, Cellpose-SAM, and user-trained U-Net models. Relevant parameters, such as expected cell diameter, flow threshold, and mask threshold, can be adjusted through the graphical user interface before processing individual files or complete directories. Following segmentation, BactoMate assigns persistent cell identifiers and calculates morphological properties for each object. These include cell area, major- and minor-axis length, aspect ratio, eccentricity, and solidity. User-defined filters can then be applied to exclude objects likely to represent debris, poorly focused cells, merged cells, or segmentation artifacts. Accepted and rejected objects are displayed in color-coded overlays, enabling users to inspect the consequences of the selected thresholds before proceeding to statistical analysis. For fluorescence images, the module calculates per-cell mean and integrated fluorescence intensities and can quantify fluorescence associated with the cell boundary. Intracellular foci can be detected using Spotiflow accompanied by a peak-detection workflow. Quantitative results are exported as structured spreadsheets together with segmentation masks, quality-control overlays, summary tables, and session parameters. Optional directory-level summaries combine measurements from batch-processed datasets.

### Cell tracking and lineage reconstruction

BactoMate integrates Trackastra for mask-based tracking of segmented cells across time-lapse image sequences. The interface provides greedy, greedy-without-division, and integer-linear-programming linking modes. A preview function applies the selected tracking parameters to an initial subset of frames 20 frames by default allowing users to inspect trajectories and optimize linking settings before processing the complete image stack. Full tracking runs generate tracked masks, trajectory tables, Cell Tracking Challenge-compatible datasets, and spreadsheet reports containing raw and filtered trajectory measurements. When division detection is enabled, BactoMate reconstructs cell lineages and generates lineage trees and founder-colored visualizations. An interactive lineage-analysis interface can be used to inspect growth, morphology, and division timing along individual pedigrees. Persistent cell identifiers connect measurements generated during segmentation and feature extraction to the corresponding trajectories and lineage assignments.

### Post-processing and batch analysis

The post-processing module combines microscopy images with quantitative measurements to generate visual outputs for inspection and presentation. These include image montages, images with calibrated scale bars, and composite videos in which time-lapse microscopy is synchronized with corresponding growth curves or other quantitative measurements. Batch processing applies a defined parameter set to multiple files within a directory. Each file is processed using the same workflow and produces its own masks, quality-control overlays, parameter records, and measurement tables. BactoMate can additionally generate directory-level summaries, allowing workflows developed for individual image sequences to be extended to experiments containing multiple strains, conditions, or fields of view without custom scripting.

### Training experiment-specific segmentation models

The training module allows segmentation models to be adapted to imaging conditions or bacterial morphologies that are not adequately represented by the supplied pretrained models. Users can construct training datasets from paired raw images and manually curated ground-truth masks and multiply the data by applying augmentation (flipping, rotation, adding noise) to increase data diversity and achieve over all better models for their specific experimental setups [24]. The module supports automatic image-mask pairing, optional cropping, and curation of training examples before model optimization. Existing Omnipose checkpoints can be fine-tuned through transfer learning, or a custom U-NET model can be trained *de novo* from user-provided annotations. Models generated by the training module can subsequently be selected in the standard segmentation workflow. Separating model development from routine analysis allows experiment-specific segmentation models to be incorporated without changing the downstream quality-control, quantification, and export procedures.

### Application of BactoMate across bacterial imaging experiments

We applied BactoMate to representative microscopy datasets spanning multichannel fluorescence imaging, bacterial motility, microcolony lineage reconstruction under phage infection conditions, and microfluidic cultivation. These datasets were assembled during a collaborative pre-release phase involving several laboratories and were selected to test whether the same graphical workflow could support distinct biological questions and hardware setups. In addition to demonstrating individual analysis functions, the applications assessed the practical transfer of workflows between experiments without requiring users to construct custom analysis scripts.

### Quantification of intracellular Ssb foci in Escherichia coli

We first analyzed multichannel instant structured illumination microscopy images of *E. coli* cells expressing the single-stranded DNA-binding (Ssb) protein fused to mNeonGreen. Cells were stained with the membrane dye FM4-64, while Ssb-mNeonGreen foci provided a fluorescent readout of intracellular single-stranded DNA formation (Fig. 3A). We then used BactoMate to convert the multichannel acquisitions, segmented individual cells using Cellpose-SAM, and detected intracellular Ssb-mNeonGreen foci using Spotiflow (Fig. 3B). Morphology-based filtering was applied to distinguish accepted cells from debris, segmentation errors, and other artifacts. BactoMate subsequently exported per-cell measurements, including the number of detected foci, cell area, and fluorescence-related parameters, as structured spreadsheets for downstream analysis and quantification (Fig. 3C). Our analysis aligns well with previous reports of SsB foci formation in non-stressed *E. coli* [25]. This application demonstrates how cell segmentation, fluorescence-foci detection, visual quality control, and quantitative export can be combined within a single workflow.

**Fig. 3.**
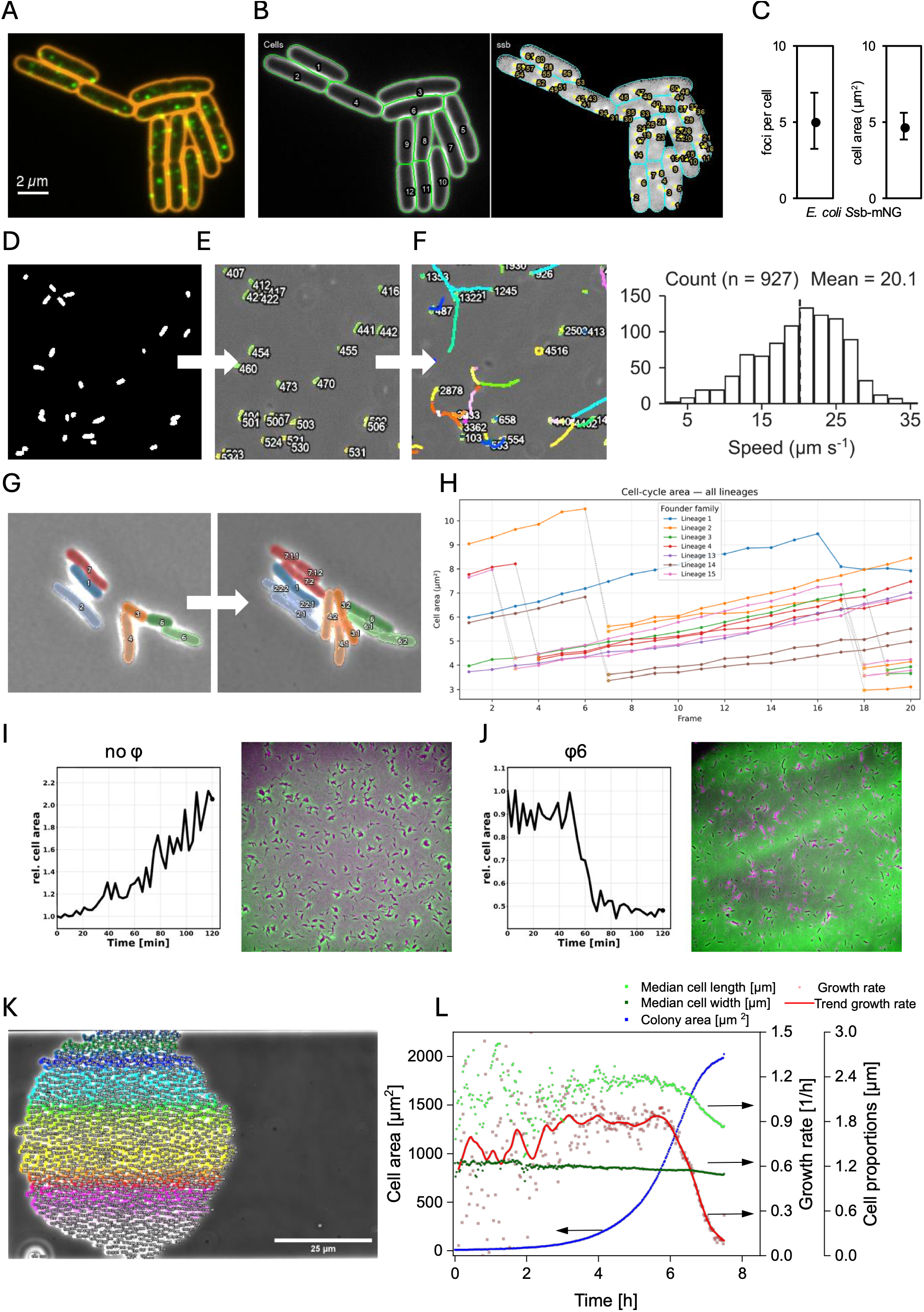
BactoMate can be used across a broad variety of microscopy experiments by expert and non-expert users. (A) Overlay of *E. coli* cells expressing Ssb-mNG and stained with a membrane dye. (B) Visual outputs generate by BactoMate of cells and foci segmentation. (C) Quantifications of foci per cell and mean cell area and standard diviation of the analyzed microscopy image. (D) Segmentation mask of swimming *Salmonella enterica* cells in liquid. (E) Preview of Trackastra within the based on the time-lapse in D (100 frames with 43ms intervalls). (F) Zoom-in of trajectories computed by Trackastra and histogram of mean speed of tracked swimming cells. (G) Trackastra tracking of cells can be also applied for lineage tracking. (H) Cell cycle of tracked lineages in G. (I, J) Time-lapse microscopy of live *Pseudomonas savastanoi* cells either uninfected (I) or infected with Φ6 at an MOI of 10 (J) and incubated at 25°C in an agar pad (overlay of phase contrast and FI images for live-death staining). Cell area changes over time were analyzed using BactoMate. (K) Colored overlay of all detected cells in the final image of the growth experiments. (L) Development of cell area, growth rate, average cell length and cell width over cultivation time.

### Tracking of swimming Salmonella enterica in liquid

To evaluate the analysis of bacterial motility, we used phase-contrast time-lapse recordings of swimming *S. enterica* cells in aqueous environment (Fig. 3D). Individual cells were segmented using Omnipose (bact_phase_omni), and trajectories were linked across consecutive frames using Trackastra. During the tracking preview, candidate trajectories were displayed directly on the microscopy images. Track-specific measurements were available through interactive tooltips, allowing users to assess whether linked detections represented continuous swimming trajectories or erroneous associations between neighboring cells (Fig. 3E). For each accepted trajectory, BactoMate calculated properties including track duration, displacement, total path length, mean speed, and straightness. The completed analysis produced a TIFF overlay displaying accepted and rejected trajectories together with a spreadsheet report containing the corresponding track-level measurements for statistical analysis (Fig. 3F).

### Reconstruction of E. coli microcolony lineages

We next investigated whether the same tracking workflow could be used to reconstruct bacterial lineages. Phase-contrast time-lapse images of *E. coli* microcolonies growing on agarose pads were converted from ND2 files to TIFF stacks while preserving pixel calibration (Fig. 3G). Images were segmented in batch mode using the Omnipose bact_phase_omni model. Morphology-based filters were applied to exclude debris, poorly segmented objects, and merged cells. The resulting masks were linked across frames using Trackastra with division detection enabled. Trajectories were initially inspected over the first 20 frames, after which the optimized settings were applied to the complete image stack. Persistent cell identifiers maintained the connection between cell morphology, trajectory identity, and lineage position. BactoMate exported per-frame summaries of cell number and total segmented area, detected division events, reconstructed lineage trees, and generated growth curves for individual lineages. Derived cell-cycle measurements were summarized in automated figures and per-lineage data tables (Fig. 3H).

### BactoMate analysis of Pseudomonas savastanoi time-lapse microscopy during φ6 infection

We next evaluated the applicability of the BactoMate workflow for analyzing bacterial time-lapse phase contrast microscopy during bacteriophage infection. To that end, we infected *P. savastanoi* with φ6, a segmented double-stranded RNA phage of the *Cystoviridae* family, at a multiplicity of infection of 10 and let the phages adsorb for 15 min. After immobilization in an agar pad, the infected cells as well as uninfected control cells were imaged every 3 min over a period of 120 min (Fig. 3I, J). We then used BactoMate to segment the image sequences in batch mode using the Cellpose-SAM model. Morphology-based filters were applied to exclude debris and poorly segmented objects, ensuring robust cell detection throughout the time course. The resulting segmentation masks were analyzed using the BactoMate post-processing video montage workflow to monitor changes in bacterial growth during infection. BactoMate exported per-frame summaries of total segmented area, detected cell division events and generated growth curves. After around 60 min of imaging (+15 min of pre-imaging adsorption), the bacterial culture collapsed in the presence of the phage, which corresponds well to the ∼75-80 min latent period of φ6 [26,27]. Thus, BactoMate provides a quantitative framework for assessing bacterial population dynamics during phage infection.

### Large-scale analysis of microfluidic growth experiments

Finally, we applied BactoMate to a high-frame-number time-lapse series of *Corynebacterium glutamicum* growing under defined substrate-limited conditions in microfluidic monolayer chambers. The dataset comprised 301 images acquired over 7.75 h and yielded more than 80,000 single-cell segmentation events. After optimization of the Omnipose parameters using the segmentation preview (see Materials and Methods), we used BactoMate to process the complete image sequence and exported temporally resolved single-cell morphology measurements and colony-level summaries (Fig. 3J). The resulting data was used to examine changes in population growth and cell morphology over time. The estimated growth rate increased modestly during the first approximately

3.5 h, remained comparatively stable for the following 2 h, and subsequently entered a period of gradual deceleration. During the transition to slower growth, median cell length decreased, whereas median cell width remained comparatively stable (Fig. 3K). This application illustrates the scalability of the BactoMate batch workflow and its ability to connect population-level growth dynamics with temporally resolved single-cell morphology.

## Conclusion

BactoMate addresses a persistent challenge in quantitative microscopy of microorganisms: the fragmentation of tools for image conversion, preprocessing, segmentation, quality control, fluorescence quantification, tracking, data export, and visualization across separate software environments. By integrating all these tasks into one user-friendly tool, BactoMate maintains parameter provenance and cell identity across successive analysis steps. This design reduces manual handoffs and makes advanced image-analysis methods accessible through a graphical interface, while the modular Python codebase preserves the possibility of extending or adapting the platform. The applications presented here illustrate the breadth of this approach. The same workflow supported multichannel fluorescence and foci quantification, tracking of free-swimming bacteria, division-aware lineage reconstruction, phage-host infection assay and analysis of a microfluidic time series. Together, these examples show that BactoMate can connect distinct imaging modalities and biological questions to a common system for quality control, single-cell measurement, and structured data export. Importantly, the examples were performed during a collaborative pre-release phase involving users from different laboratories, indicating that the platform can be rapidly adopted outside its immediate development environment.

Parameter transparency is a central design principle of BactoMate. User-defined settings are exposed in the interface and stored in session-specific JSON records together with the corresponding outputs. These records facilitate the consistent reuse of analysis settings across biological replicates and provide a traceable link between quantitative results and the parameters used to generate them. Visual overlays of accepted and rejected objects further allow users to inspect the consequences of morphology-based quality-control thresholds before population statistics are calculated. This is particularly important when biological conclusions depend on comparatively small differences in properties such as cell width, fluorescence intensity, or division timing. However, parameter provenance alone does not guarantee complete computational reproducibility. Exact reproduction also depends on software and model versions, dependency environments, and, where applicable, hardware-dependent execution. BactoMate provides flexible segmentation and tracking workflows that can be adapted to diverse imaging conditions, cell densities, bacterial morphologies, and experimental designs. The supplied U-Net models support a broad range of bacterial datasets, while adjustable parameters and the modular architecture enable users to optimize existing models, apply transfer learning, or train custom models for specialized applications. Tracking similarly benefits from configurable workflows for different temporal resolutions, motility patterns, and division dynamics. Interactive previews and visual quality-control tools allow users to assess segmentation and tracking results directly and refine the analysis before quantitative measurements are generated. Together, these features support the development and validation of analysis configurations tailored to each experimental system, while preserving a consistent workflow for single-cell quantification and data export.

BactoMate is distributed as open-source software under the MIT license through our public GitHub repository and is accompanied by a series of tutorial videos on YouTube to facilitate installation, onboarding, and adoption by the broader community. Its modular architecture provides a foundation for incorporating additional segmentation and tracking backends, quantitative descriptors, file formats, and visualization methods as bacterial imaging experiments increase in scale and complexity. Future developments could include automated quality metrics, improved workflow and model versioning, greater interoperability with community image-data standards, and mechanisms for sharing validated analysis configurations. Through continued benchmarking and community-driven development, BactoMate has the potential to evolve from an integrated desktop analysis environment into a reusable framework for transparent, reproducible, and comparable microbial single-cell microscopy workflows.

## Methods

### Software implementation

BactoMate is implemented in Python 3.11 with a PyQt6 graphical interface. Core dependencies include Omnipose, cellpose-omni, Cellpose-SAM, Spotiflow, Trackastra, bioio, nd2, readlif, czifile, NumPy, SciPy, scikit-image, pandas, and PyTorch. The codebase is organized into GUI modules (conversion, segmentation, analysis, tracking, post-processing, training) and shared libraries for mask I/O, morphology metrics, foci detection, cell-ID linking, and visualization. The U-NET training module was adapted from bnsreenu Unet_nuclei_tutorial (uploaded: 2020; accessed Dezember 2025) (https://github.com/bnsreenu/python_for_microscopists/blob/master/076-077-078-Unet_nuclei_tutorial.py). Long-running operations execute in background worker threads. Compatibility patches for Omnipose and cellpose-omni are applied at installation. Continuous integration runs installation, smoke tests, and pytest on macOS and Windows.

### Procedure of presented example microscopy experiments

#### iSIM acquisition of Ssb-mNG expression in E. coli cells

*E. coli* strains were grown in LB at 30 °C to mid-exponential phase and prepared on agarose pads for time-lapse imaging. Multi-channel fluorescence experiments used FM4-64 membrane staining and Ssb-mNeonGreen replication focus reporters as described previously. Imaging was performed using a VisiTech VT-iSIM Super Resolution Imaging System configured for instant structured illumination microscopy (iSIM) on a Nikon Ti2-E inverted microscope, equipped with a Nikon CFI SR Apochromat TIRF 100× oil-immersion objective (NA 1.49, working distance 0.12 mm) and dual Hamamatsu ORCA-Quest C15550-20UP qCMOS cameras with 4.6 × 4.6 µm sensor pixels; the pixel size was stored in the acquisition metadata and propagated by BactoMate during conversion.

#### Single-cell swimming assay

Data was re-analyzed from [28]. In brief, the experiment was performed as follows. Overnight cultures were incubated in LB at 30°C and180rpm. Subcultures were inoculated1:100 in 10 mL fresh LB and cultivated accordingly. After 2.5 h of growth, *flhDC* expression was synchronized by induction with AnTc, (final concentration= 100 ng/ml) followed by 30 min of incubation. Subsequently, cells were harvested at 2,500×g for 5 min and resuspended in fresh AnTc-free media. Directly, at 0 min post medium switch a first sample was drawn and processed for obtaining swimming behavior. For this, 70 μL were loaded in a flow cell and microscopy was performed using a Ti-2 Nikon inverted microscope equipped with a CFI Plan Apochromat DM 20×Ph2/0.75 objective. Data was acquired for 100 frames with a time interval of 43 ms between frames.

#### Single-cell growth assay lineage tracking

Data was re-analyzed from [29]. In brief, the experiment was performed as follows. Overnight cultures of *E. coli* strains carrying EcDruIII plasmids were grown with shaking at 180 rpm in LB Lennox medium containing 10 mM MgSO_4_ and 2 mM CaCl_2_, supplemented with 100 µg ml^-1^ ampicillin at 30 °C. The following day, a 1:100 subculture was inoculated and grown at 30 °C until an OD_600_ of 0.3–0.5 was reached. Subsequently, 1 µl of the culture was spotted onto a 1.2% (w/v) agarose pad (prepared in LB:MQ at a 1:5 ratio). Microscopy slides were mounted in an incubation chamber preheated to 30 °C. Image acquisition was performed using a Nikon Eclipse Ti2 inverted microscope equipped with a CFI Plan Apochromat DM ×60 Lambda oil Ph3/1.40 objective. Phase-contrast images were captured every 2 min for 20 frames.

#### Time-lapse microscopy of Pseudomonas savastanoi in the presence and absence of phage

An overnight culture of *Pseudomonas savastanoi* LM2489 was diluted 1:100 and grown at 25°C with shaking at 200 rpm to an OD_600_ of 0.3. Cells were infected with Φ6 at an MOI of 10 for 15 min at 25°C in a total volume of 500 µl to allow adsorption. Following adsorption, cells were pelleted by centrifugation at 5000 rcf for 5 min, the supernatant discarded, and the pellet resuspended in 50 µl of 1x PBS supplemented with 0.2 µg/ml propidium iodide (PI). Cells were then immobilized in a 1% agarose pad prepared in 20% LB in 1x PBS. Time-lapse imaging was performed using an epifluorescence microscope (Leica) equipped with an LED light engine for fluorescence illumination and a DFC9000GT-VSC11903 camera. Cells were maintained at 25°C using an Incubator i8 (PECON) during imaging. Images were acquired every 3 min for 120 min using a 40x air objective.

#### Microfluidic chip preparation, cultivation, microscopy setup and BactoMate analysis

The picoliter batch reactor was designed and fabricated as previously described by Smaluch et al. [30]. Pre-cultivation of *Corynebacterium glutamicum* WT ATCC 13032 was performed in two sequential steps. First, a 100 mL baffled shake flask containing 20 mL brain-heart-infusion (BHI) complex medium was inoculated from ROTI^®^Store cryo-vials (Carl Roth, Germany). Following overnight incubation at 30 °C and 120 rpm, a second pre-culture was inoculated to an initial optical density OD_600_ of 0.2. This culture was incubated under identical conditions for 3 hours. During their exponential growth phase, the cells were harvested via centrifugation (5000 rpm, 5 mins) and resuspended in fresh BHI medium to a final OD_600_ of 0.1 for subsequent microfluidic cultivation. Microscopy was performed using a Nikon Eclipse Ti2 microscope, equipped with a 100x oil immersion objective (Nikon Corporation, Japan). The microfluidic cultivation device was mounted on a motorized stage and surrounded by a microscope incubator to cultivate at a constant 30°C. Inoculation procedure was performed as described in [30]. Phase contrast images were acquired every 1.5 min for 7.75 h with a 1T-01-N-KINETIX-M-C camera (Teledyne Photometrics, USA), resulting in 301 frames. For image analysis with

#### BactoMate analysis parameters

Analyses were performed with BactoMate v2.8.4 or subsequent versions. Segmentation used 4_phase_omni (Omnipose) or Cellpose-SAM with diameter 30 px, flow threshold 0.4, GPU enabled. Morphology filters: area, aspect ratio, and eccentricity thresholds as indicated in session logs. Foci detection used Spotiflow with user-defined score and overlap thresholds. Tracking used Trackastra with greedy linking and division detection. Batch processing was applied across all fields of view. Exported Excel and CSV files were analyzed for population statistics. For the microfludic experiment derrived data in BactoMate the image series was cropped, rotated and aligned by using Fiji (Version 1.54) [31]. For segmentation with BactoMate the “bacto_phase_omni” model was selected. The preview segmentation windows were used to change the cell segmentation parameters to 1.00 for flow and 0.2 for mask. This pixel size was adjusted to 0.065 µm. The CSV datafiles generated by BactoMate were used for analysis of the growth rate according to [32]. Graphs were generated using OriginPro 2023. Growth rate trend was calculated using the Lowess filter provided by Origin. Median cell length and width were estimated using the descriptive statistic function.

#### Online tutorial

We provide a set of video tutorials introducing BactoMate, aiding installation procedure and applying BactoMate for a set of microscopy analyses via our publicly available YouTube channel: https://www.youtube.com/playlist?list=PLYQZb_PwGpYw.

## Code availability

BactoMate is open-source software (MIT license). Source code, install scripts, and documentation are available at https://github.com/Molinfect/BactoMate. BactoMate is installable via a stand-alone dmg file for macOS and per setup/install_windows.bat for Windows. User documentation is provided in docs/BACTOMATE_USER_MANUAL.md.

## Acknowledgements

We thank all members of the Erhardt and Popp laboratories for testing BactoMate and providing feedback. We acknowledge the developers of Omnipose, Cellpose, Spotiflow, and Trackastra, whose open-source tools form the analytical backbone of BactoMate. P.F.P and M.E. acknowledge funding from the Deutsche Forschungsgemeinschaft (DFG) in the framework of the priority program SPP2330 (research grant no. 548567920 to P.F.P (PO 2831/2-1) and to M.E. (ER 778/13-1)). The VT-iSIM Super Resolution Imaging System was funded by the DFG, project number 545038525 (INST 276/868-1 FUGG). MP and DK acknowledge funding from the Deutsche Forschungsgemeinschaft (DFG) in the framework of the priority program SPP2170. This work was further supported by the DFG-funded SPP2389 (grant number 569011257 to J.H.).

## Author contributions

P.F.P. conceived the project and together with L.H. and S.F. wrote the software. P.F.P. together with M.P. and S. A. performed imaging experiments and validated workflows. M.E., P.F.P., D. K. and J. H. supervised the project. P.F.P. wrote the manuscript with input from all authors.

